# Analyzing the impact of SARS CoV-2 on the human proteome

**DOI:** 10.1101/2020.05.21.107912

**Authors:** Ernesto Estrada

## Abstract

The COVID-19 respiratory disease is caused by the novel coronavirus SARS-CoV-2, which uses the enzyme ACE2 to entry human cells. This disease is characterized by important damages at multi-organ level, partially due to the abundant expression of ACE2 in practically all human tissues. However, not every organ in which ACE2 is abundant is affected by SARS CoV-2, which suggests the existence of other multi-organ routes for transmitting the perturbations produced by the virus. We consider here diffusive processes through the protein-protein interaction (PPI) network of proteins targeted by SARS CoV-2 as such alternative route. We found a subdiffusive regime that allows the propagation of virus perturbations through the PPI network at a significant rate. By following the main subdiffusive routes across the PPI network we identify proteins mainly expressed in the heart, cerebral cortex, thymus, testis, lymph node, kidney, among others of the organs reported to be affected by COVID-19.

## 1 Introduction

Since December 2019 a new coronavirus designated SARS-CoV-2 (severe acute respiratory syndrome coronavirus 2) (*1*) has produced an outbreak of pulmonary disease which has soon became a global pandemic (*2, 3*). The new coronavirus disease (COVID-19) is characterized by a wide range of clinical manifestations (*4, 5*), with an important implication of multi-organ and systemic dysfunctions (*6, 7*), which include heart failure, renal failure, liver damage, shock and multi-organ failure. The new coronavirus shares about 82% of its genome with the one which produced the 2003 outbreak (SARS CoV) (*8*). Both coronaviruses share the same cellular receptor, which is the angiotensin-converting enzyme 2 (ACE2) receptor (*8, 9, 10*). ACE2 receptors are enriched in several tissues across the organs, such as in alveolar epithelial type II cells of lung tissues, in the heart, endothelium, kidneys and intestines. Therefore, ACE2 has been hypothesized as a potential cause of the major complications of the COVID-19 (*11, 12*). However, it has been found that ACE2 has abundant expression on endothelia and smooth muscle cells of virtually all organs (*13*). Therefore, it should be expected that after SARS CoV-2 is present in circulation, it can be spread across all organs. In contrast, both SARS CoV and SARS CoV-2 are found specifically in some organs but not in others, as shown by *in situ* hybridization studies for SARS CoV [13]. This was already remarked by Hamming et al. (*13*) by stressing that it “*is remarkable that so few organs become viruspositive, despite the presence of ACE2 on the endothelia of all organs and SARS-CoV in blood plasma of infected individuals”.*

Recently Gordon et al. (*14*) identified human proteins that interact physically with those of the SARS CoV-2 forming a high confidence SARS-CoV-2-human protein-protein interactions (PPIs) system. Using this information Gysi et al. (*15*) discovered that 208 of the human proteins targeted by SARS CoV-2 forms a connected component inside the human PPI network. That is, these 208 are not randomly distributed across the human proteome but they are closely interconnected by short routes that allow moving from one to another in just a few steps. These interdependencies of protein-protein interactions are known to enable that perturbations on one interaction propagates across the network and affect other interactions (*16, 17, 18, 19*). In fact, it has been signified that diseases are a consequence of such perturbation propagation (*20, 21, 22*). It has been stressed that the protein-protein interaction process requires diffusion in their initial stages (*23*). The diffusive processes occur when proteins, possibly guided by electrostatic interactions, need to encounter each other many times before forming an intermediate (*24*). Not surprisingly, diffusive processes have guided several biologically oriented searches in PPI networks (*25, 26*). Therefore, we assume here that perturbations produced by SARS CoV-2 proteins on human PPI network are propagated by means of diffusive processes. However, due to the crowded nature of the intra-cell space and the presence in it of spatial barriers, subdiffusive processes more than normal diffusion are expected for these protein-protein encounters (*27, 28, 29*). This creates another difficulty, as remarked by Batada et al. (*23*), which is that such (sub)diffusive processes along are not sufficient for carrying out cellular processes at a significant rate in cells.

Here we propose the use of a time-fractional diffusion model on the PPI network of proteins targeted by SARS CoV-2. The goal is to model the propagation of the perturbations produced by the interactions of human proteins with those of SARS CoV-2 through the whole PPI. The subdiffusive process emerging from the application of this model to the SARS-CoV-2-human PPIs has a very small rate of convergence to the steady state. However, this process produces dramatic increment of the probability that certain proteins are perturbed at very short times. This kind of shock wave effect of the transmission of perturbations occurs at much earlier times in the subdiffusive regime than at the normal diffusion one. Therefore, we propose here a switch and restart process in which a subdiffusive process starts at a given protein of the PPI, perturbs a few others, which then become the starting point of a new subdiffusive processm and so on. Using this approach we then analyze how the initial interaction of the SARS CoV-2 Spike protein with a human protein propagates across the whole network. We discover some potential routes of propagation of these perturbations from proteins mainly expressed in the lungs to proteins mainly expressed in other different tissues, such as the heart, cerebral cortex, thymus, lymph node, testis, prostate, liver, small intestine, duodenum, kidney, among others.

## 2 Materials and Methods

### 2.1 Time-fractional diffusion model on networks

In this work we always consider Γ = (*V, E*) to be an undirected finite network with vertices *V* representing proteins and edges E which represent the interaction between pairs of proteins. Let us consider 0 < *α* ≤ 1 and a function 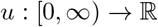, then we denote by 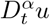 the fractional Caputo derivative of *u* of order *α*, which is given by (*31*)

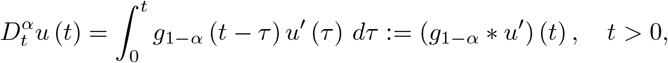

where ∗ denotes the classical convolution product on (0, ∞) and 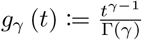, for γ > 0. Observe that the previous fractional derivative has sense whenever the function is derivable and the convolution is defined (for example if u’ is locally integrable). The notation *g*_γ_ is very useful in the fractional calculus theory, mainly by the property *g*_γ_ ∗ *g_δ_* = *g*_γ+*δ*_ for all *γ, δ* > 0.

Here we propose to consider the time-fractional diffusion (TFD) equation on the network as

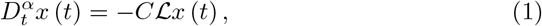

with initial condition *x* (0) = *x*_0_, where *x_i_* (*t*) is the probability that protein *i* is perturbed at time *t, C* is the diffusion coefficient of the network, which we will set hereafter to unity, and 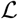 is the graph Laplacian, i.e., 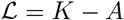, where *K* is a diagonal matrix of node degrees and *A* is the adjacency matrix. This model was previously studied in distributed coordination algorithms for the consensus of multi-agent systems (*32, 33, 34*). The use of fractional calculus in the context of physical anomalous diffusion has been reviewed by Metzler and Klafter (*30*). The solution of this TFD equation is given by:

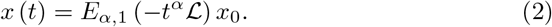

where *E_α,β_* (·) is the Mittag-Leffler matrix function. This result is proved in the Supplementary Note 1.

We can write 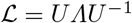, where 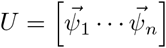 and *Λ = diag* (*μ_r_*). Then,

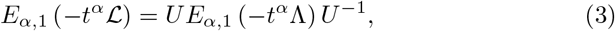

which can be expanded as

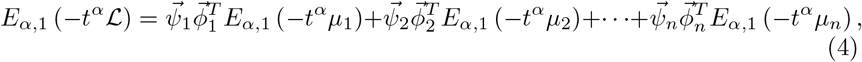

where 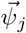 and 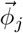 are the *j*th column of *U* and of *U*^−1^, respectively. Because *μ*_1_ = 0 and 0 < *μ*_2_ ≤ ⋯ ≤ *μ_n_* for a connected graph we have

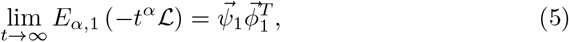

where 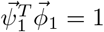. Let us take 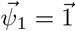, such that we have

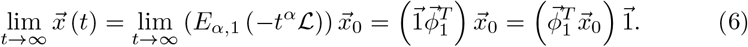

This result indicates that in an undirected and connected network, the diffusive process controlled by the TFD equation always reaches a steady state which consists of the average of the values of the initial condition. Also, because the network is connected *μ*_2_ makes the largest contribution to 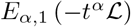 among all the nontrivial eigenvalues of 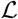. Therefore, it dictates the rate of convergence of the diffusion process. We remark that in practice the steady state lim_*t*→∞_ ||*x_v_* (*t*) − *x_w_* (*t*)|| = 0, ∀*v, w* ∈ *V* is very difficult to achieve. Therefore, we use a threshold *ε*, e.g., *ε* = 10^−3^, such that lim_*t*→∞_ ||*x_v_* (*t*) − *x_w_* (*t*)|| = *ε* is achieved in relatively small simulation time.

Due to its importance in this work we remark the structural meaning of the Mittag-Leffler function of the Laplacian matrix appearing in the solution of the TFD equation. That is, 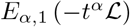 is a matrix function, which is defined as

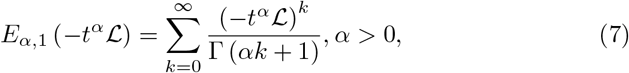

where Γ (·) is the Euler gamma function. We remark that for *α* =1 we recover the diffusion equation on the network: 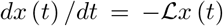 and its solution 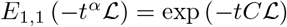 is the well-known heat kernel of the graph.

### 2.2 Time-fractional diffusion distance

We define here a generalization of the diffusion distance studied by Coifman and Lafon (*35*). We start by defining

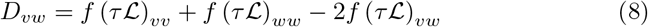

where 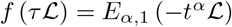. In the Supplementary Note 2 we prove that *D_vw_* is a square Euclidean distance between the nodes *v* and *w*. Let *μ_j_* be the *j*th eigenvalue and *ψ_ju_* the uth entry of the *j*th eigenvector of the Laplacian matrix. Then, we can write the time-fractional diffusion distance as

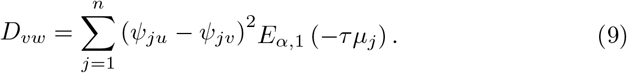

It is evident that when *α* = 1, *D_vw_* is exactly the diffusion distance previously studied by Coifman and Lafon (*35*). The fractional-time diffusion distance between every pair of nodes in a network can be represented in a matrix form as follow:

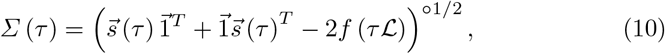

where 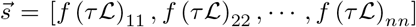 is a vector whose entries are the main diagonal terms of the Mittag-Leffler matrix function, 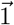 is an all-ones vector and ○ indicates an entrywise operation. Using this matrix we can build the diffusion distance-weighted adjacency matrix of the network:

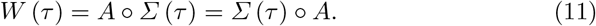

The shortest diffusion path between two nodes is then the shortest weighted path in *W* (*τ*). Let us consider each of the terms forming the definition of the time-fractional diffusion distance and apply the limit of the very small −*t^α^*. That is,

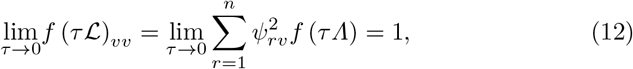

and in a similar way,

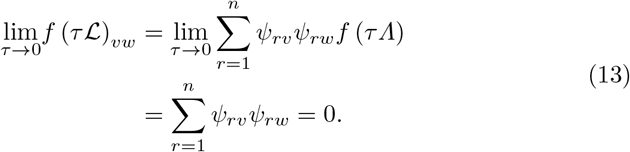

Therefore, 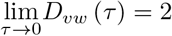. Consequently, 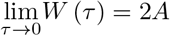, which immediately implies that the time-fractional shortest diffusion path is identical to the shortest (topological) one in the limit of very small *τ* = − *t^α^*.

### 2.3 Network of proteins targeted by SARS CoV-2

The proteins of SARS CoV-2 and their interactions with human proteins were determined experimentally by Gordon et al. (*14*). Gysi et al. (*15*) constructed an interaction network of all 239 human proteins targeted by SARS CoV-2. In this network the nodes represent human proteins targeted by SARS CoV-2 and two nodes are connected if the corresponding proteins have been determined to interact with each other. Obviously, this network of proteins targeted by SARS CoV-2 is a subgraph of the protein-protein interaction (PPI) network of humans. One of the surprising findings of Gysi et al. (*15*) is the fact that this subgraph is not formed by proteins randomly distributed across the human PPI, but they form a main cluster of 208 proteins and a few small isolated components. Hereafter, we will always consider this connected component of human proteins targeted by SARS CoV-2. This network is formed by 193 proteins which are significantly expressed in the lungs. Gysi et al. (*15*) reported a protein as being significantly expressed in the lungs if its GTEx median value is larger than 5. GTEx (*36*) is a database containing the median gene expression from RNA-seq in different tissues. The other 15 proteins are mainly expressed in other tissues. However, in reporting here the tissues were proteins are mainly expressed we use the information reported in The Human Protein Atlas (*37*) were we use information not only from GTEx but also from HPA (see details at the Human Protein Atlas webpage) and FANTOM5 (*38*) datasets.

## 3 Results

### 3.1 Global characteristics of the fractional diffusion

The PPI network of human proteins targeted by SARS CoV-2 is very sparse, having 360 edges, i.e., its edge density is 0.0167, 30% of nodes have degree (number of connections per protein) equal to one and the maximum degree of a protein is 14. The second smallest eigenvalue of the Laplacian matrix of this network is very small, i.e., *μ*_2_ = 0.0647. Therefore, the rate of convergence to the steady state of the diffusion processes taking place on this PPI are very slow. We start by analyzing the effects of the fractional coefficient *α* on these diffusive dynamics. We use the normal diffusion *α* = 1 as the reference system.

To analyze the effects of changing *α* over the diffusive dynamics on the PPI network we consider the solution of the TFD equation for processes starting at a protein with a large degree, i.e., PRKACA, degree 14, and a protein with a low degree, i.e., MRPS5, degree 3. That is, the initial condition vector consists of a vector having one at the entry corresponding to either PRKACA or MRPS5 ans zeroes elsewhere. In Fig. 1 we display the changes of the probability with the shortest path distance from the protein where the process starts. This distance corresponds to the number of steps that the perturbation needs to traverse to visit other proteins. For *α* = 1.0 the shape of the curves in Fig. 1 are the characteristic ones for the Gaussian decay of the probability with distance. However, for *α* < 1 we observe that such decay differs from that typical shape showing a faster initial decay followed by a slower one. In order to observe this effect in a better way we zoomed the region of distances from 2 to 4 (see 1 (b) and (d)). As can be seen for distances below 3 the curve for *α* = 1.0 is on top of those for *α* < 1 indicating a slower decay of the probability. After this distance, there is an inversion and the normal diffusion occurs at much faster rate than the other two for the longer distances. This is a characteristic signature of subdiffusive processes, which starts at much faster rates than a normal diffusive process and then continue at much slower rates. Therefore, here we observe that the subdiffusive dynamics are much faster at earlier times of the process, which is when the perturbation occurs to close nearest neighbors to the initial point of perturbation.

**Fig. 1:**
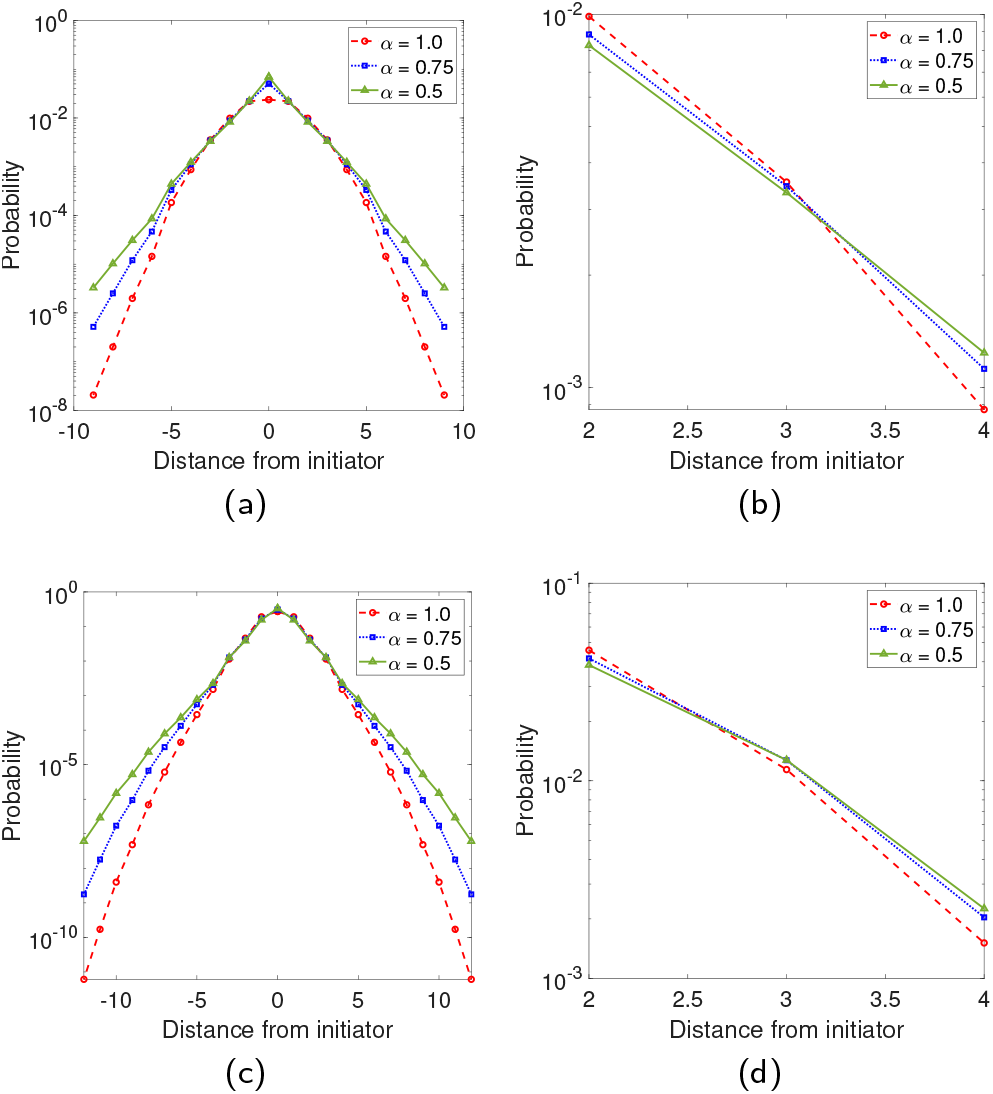
Spatial propagation of perturbations. Average probability that a protein at a given distance from the perturbed protein feels the perturbation. (A) The initiators are the protein PRKACA (a) and (b) as well as MRPS5 (c) and (d). (b) and (d) zoom the scale of the processes in (A) and (C), respectively.

To further investigate these characteristic effects of the subdiffusive dynamics we study the time evolution of a perturbation occurring at a given protein and its propagation across the whole PPI network. In Fig. 2 we illustrate these results for *α* = 1.0 (a), *α* = 0.75 (a), *α* = 0.5 (c). As can be seen in the main plots of this Figure the rate of convergence of the processes to the steady state is much faster in the normal diffusion (a) than in the subdiffusive one (b) and (c). However, at very earlier times (see insets in Fig. 2) there is a shock wave increase of the perturbation at a set of nodes. Such kind of shock waves have been previously analyzed in other contexts as a way of propagating effects across PPI networks (*17*). The remarkable finding here is however the fact that such shock wave occurs at much earlier times in the subdiffusive regimes than at the normal diffusion. That is, while for *α* = 1.0 these perturbations occur at *t* ≈ 0.1 − 0.3, for *α* = 0.75 they occur at *t* ≈ 0.0 − 0.2, and for *α* = 0.5 at *t* ≈ 0.0 − 0.1. Seeing this phenomenon in the light of what we have observed in the previous paragraph is not strange due to the observation that such processes go at much faster rate at earlier times, and at short distances, than the normal diffusion. In fact, this is a consequence of the existence of a positive scalar *T* for which *E*_*α*,1_ (− *γt^α^*) decreases faster than exp(− *γt*) for *t* ∈ (0, *T*) for 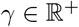 and 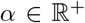 (see Theorem 4.1 in (*32*)). Hereafter, we will consider the value of *α* = 0.75 for our experiments due to the fact that it reveals a subdiffusive regime, but the shoch waves observed before are not occurring in an almost instantaneous way like when *α* = 0.5, which would be difficult from a biological perspective.

**Fig. 2:**
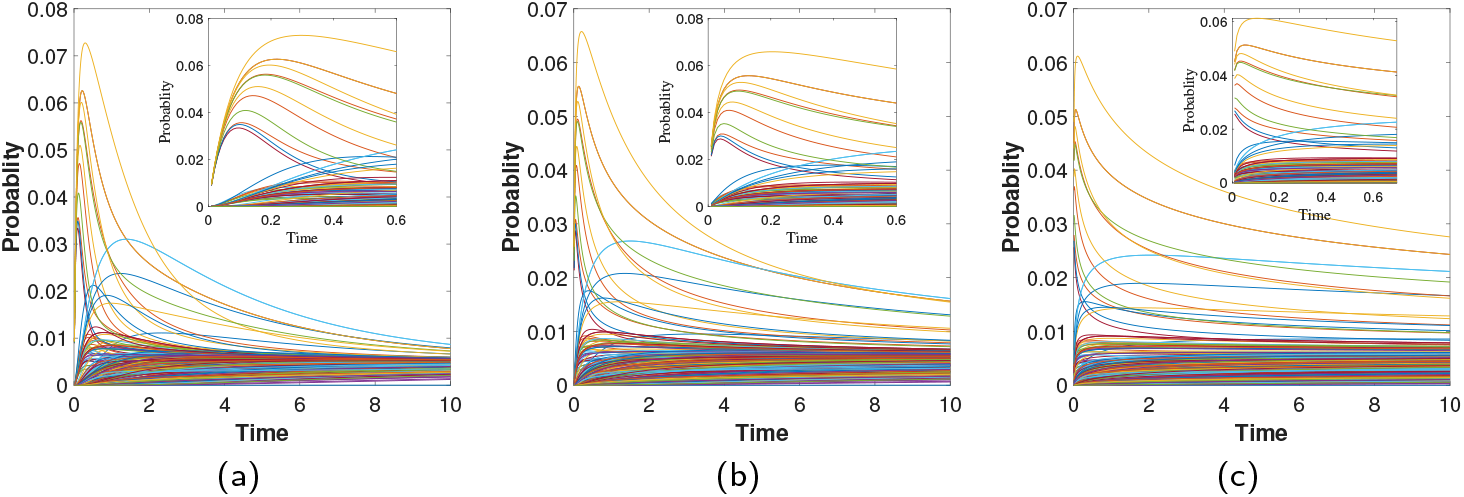
Shock wave increase of perturbations across the network. Time evolution of the propagation of perturbations from the protein PRKACA to the rest of the proteins in the PPI network of human proteins targeted by SARS CoV-2. (a) Normal diffusion *α* = 1.0. (b) Subdiffusion obtained for *α* = 3/4. (c) Subdiffusion obtained for *α* = 1/2. The insets illustrate the shortest time evolution of the perturbations.

The previous results put us in a crossroad. First, the subdiffusive processes which are expected due to the crowded nature of the intra-cellular space, are very slow for carrying out cellular processes at a significant rate in cells. However, the perturbation shocks occurring at earlier times of these processes are significantly faster than in normal diffusion. To sort out these difficulties we propose a switching back and restart subdiffusive process occurring in the PPI network. That is, a subdiffusive process starts at a give protein, which is directly perturbed by a protein of SARS CoV-2. It produces a shock wave increase of the perturbation in close neighbors of that proteins. Then, a second subdiffusive process starts at these newly perturbed proteins, which will perturb their nearest neighbors. The process is repeated until the whole PPI network is perturbed. This kind of “switch and restart processes” have been proposed for engineering consensus protocols in multiagent systems (*40*) as a way to accelerate the algorithms using subdiffusive regimes.

### 3.2 Diffusion from SARS CoV-2 Spike protein

The so-called Spike protein (S-protein) of the SARS CoV-2 interacts with only two proteins in the human hosts, namely ZDHHC5 and GOLGA7. The first protein, ZDHHC5, is not in the main connected component of the PPI network of SARS CoV-2 targets. Therefore, we will consider here how a perturbation produced by the interaction of the virus S-protein with GOLGA7 is propagated through the whole PPI network of SARS CoV-2 targets. GOLGA7 has degree one in this network and its diffusion is mainly to close neighbors, namely to proteins separated by 2-3 edges. When starting the diffusion process at the protein GOLGA7 the main increase in the probability of perturbing another protein is reached for protein GOLGA3, which increases its probability up to 0.15 at *t* = 0.2, followed by PRKAR2A, with a small increase in its probability, 0.0081. Then, the process switch and restarts at GOLGA3, which mainly triggers the probability of the protein PRKAR2A–a major hub of the network. Once we start the process at PRKAR2A, practically the whole network is perturbed with probabilities larger than 0.1 for 19 proteins apart from GOLGA3. These proteins are, in decreasing order of their probability of being perturbed: AKAP8, PRKAR2B, CEP350, MIB1, CDK5RAP2, CEP135, AKAP9, CEP250, PCNT, CEP43, PDE4DIP, PRKACA, TUB6CP3, TUB6CP2, CEP68, CLIP4, CNTRL, PLEKHA5, and NINL. Notice that the number of proteins perturbed is significantly larger than the degree of the activator, indicating that not only nearest neighbors are activated.

An important criterion for revealing the important role of the protein PRKAR2A as a main propagator in the network of proteins targeted by SARS CoV-2 is its average diffusion path length. This is the average number of steps that a diffusive process starting at this protein needs to perturb all the proteins in the network. We have calculated this number to be 3.6250, which is only slightly larger than the average (topological) path length, which is 3.5673. That is, in least than 4 steps the whole network of proteins is activated by a diffusive process starting at PRKAR2A. Also remarkable, the average shortest diffusive path length is almost identical to the shortest (topological) one. This means that this protein mainly uses shortest (topological) paths in perturbing other proteins in the PPI. In other words, it is highly efficient in conducting such perturbations. We will analyze this characteristics of the PPI of human proteins targeted by SARS CoV-2 in a further section of this work.

At this time almost any protein in the PPI network is already perturbed. Therefore we can switch and restart the subdiffusion from practically any protein at the PPI network. We then investigate which are the proteins with the higher capacity of activating other proteins which are involved in human diseases. Here we use the database SiGeNet (*39*), which is one of the largest publicly available collections of genes and variants associated to human diseases. We identified 38 proteins targeted by SARS CoV-2 for which there are “definitive” or “strong” evidence of being involved in a human disease or syndrome (see Supplementary Table 1). These proteins participate in 70 different human diseases or syndromes as given in Supplementary Tables 2 and 3. We performed an analysis in which a diffusive process starts at any protein of the network and we calculated the average probability that all the proteins involved in human diseases are then perturbed. For instance, for a subdiffusive process starting at the protein ARF6 we summed the probabilities that the 38 proteins involved in diseases are perturbed at an early time of the process *t* = 0.2. Then, we obtain a global perturbation probability of 0.874. By repeating this process for every protein as an initiator we obtained the top disease activators. We have found that none of the 20 top activators is involved itself in any of the human diseases or syndromes considered here. They are, however, proteins which are important not because of their direct involvement in diseases or syndromes but because they propagate perturbations in a very effective way to those directly involved in such diseases/syndromes. Among the top activators we have found: ARF6, ECSIT, RETREG3, STOM, HDAC2, EXOSC5, THTPA, among others shown in Fig. 4, where we illustrate the PPI network of the proteins targeted by SARS CoV-2 remarking the top 20 disease activators.

**Fig. 4:**
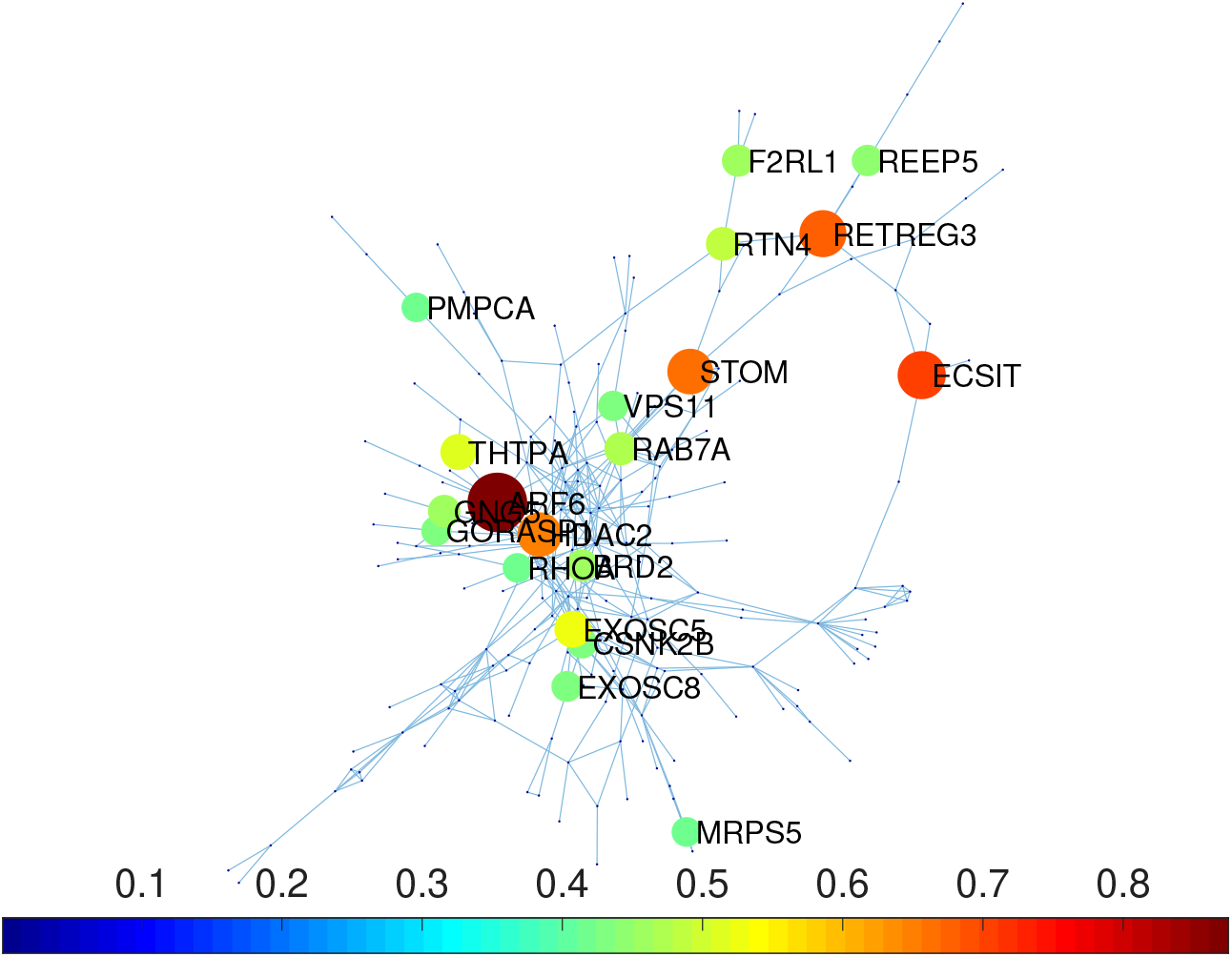
Main disease activators. Proteins targeted by SARS CoV-2 identified as the top 20 main activators of proteins which are involved in human diseases.

### 3.3 Diffusion from the lungs to other organs

We now consider how a perturbation produced by SARS CoV-2 on a protein mainly expressed in the lungs can be propagated to proteins mainly located in other tissues (see Supplementary Table 4) by a subdiffusive process. That is, we start the subdiffusive process by perturbing a given protein which is mainly expressed in the lungs. Then, we observe the evolution of the perturbation at every of the proteins mainly expressed in other tissues. We repeat this process for all the 193 proteins mainly expressed in the lungs. In every case we record those proteins outside the lungs which are perturbed at very early times of the subdiffusive process. For instance, in Fig. 3 we illustrate one example in which the initiator is the protein GOLGA2, which triggers a shock wave on proteins RBM41, TL5 and PKP2, that are expressed mainly outside the lungs. We consider such perturbations only if they occur at *t* < 1.

**Fig. 3:**
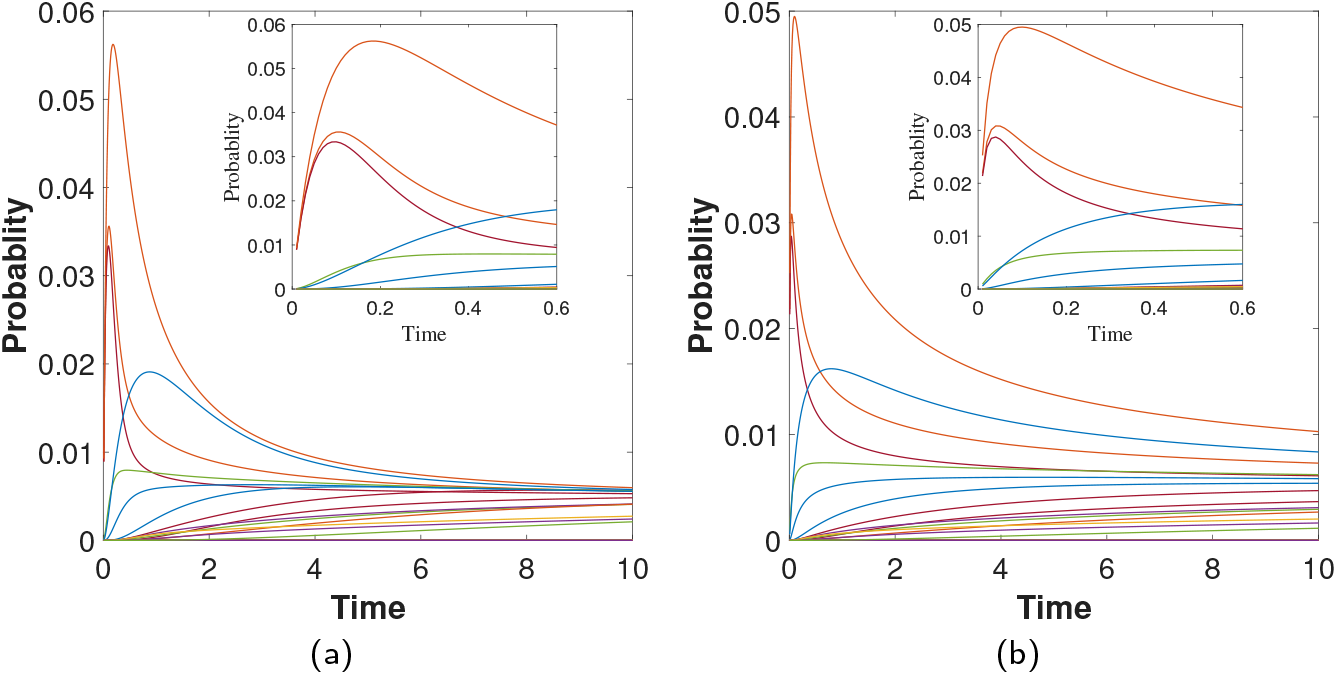
Shock wave increase of perturbations in proteins outside the lungs. Time evolution of the propagation of perturbations from the protein GOLGA2 to proteins mainly expressed outside the lungs. (a) Normal diffusion *α* = 1.0. (b) Subdiffusion with *α* = 3/4. The insets illustrate the shortest time evolution of the perturbation.

Not every one of the proteins expressed outside the lungs is triggered by such shock waves at very early time of the diffusion. For instance, proteins MARK1 and SLC27A2are perturbed in very slow processes, and do not produce the characteristics high peaks in the probability at very short times. On the other hand, there are proteins expressed outside the lungs which are triggered by more than one protein from the lungs. The case of GOLGA2 is an example of a protein triggered by 3 ones in the lungs. In Table 1 we list some of the proteins expressed mainly in tissues outside the lungs which are heavily perturbed by proteins in the lungs. The complete list of the perturbing proteins is given in the Supplementary Table 5. We give three indicators of the importance of the perturbation of these proteins. They are: Act., which is the number of proteins in the lungs that activate each of them; *p*_tot_, which is the sum of the probabilities of finding the diffusive particle at this protein for diffusive processes that have started in the their activators; *t*_mean_, which is the average time required by activators to perturb the corresponding protein. For instance, PKP2 is perturbed by 21 proteins in the lungs, which indicates that this protein, mainly expressed in the heart muscle, has a large chance of being perturbed by diffusive processes starting in proteins mainly located at the lungs. Protein PRIM2 is activated by 5 proteins in the lungs, but if all these proteins were acting at the same time, the probability that PRIM2 is perturbed will be very high, *p*_tot_ ≈ 0.536. Finally, protein TLE5 is perturbed by 13 proteins in the lungs, which needs as average *t*_mean_ ≈ 0.24 to perturb TLE5. These proteins do not form a connected component among them in the network. The average shortest diffusion path between them is 5.286 with a maximum shortest subdiffusion path of 10. As average they are almost equidistant from the rest of the proteins in the network as among themselves. That is, the average shortest subdiffusion path between these proteins expressed outside the lungs and the rest of the proteins in the network is 5.106. Therefore, these proteins can be reached from other proteins outside the lungs in no more than 6 steps in subdiffusive processes like the ones considered here.

**Tab. 1:** Multi-organ propagation of perturbations. Protein mainly expressed outside the lungs which are significantly perturbed during diffusive processes started at other proteins expressed in the lungs. Act., is the number of lungs proteins activators; *p*_tot_, is the sum of the probabilities of finding the diffusive particle at this protein; *t*_mean_, is the average time of activation (see text for explanations). The tissues of main expression are selected among the ones with the highest Consensus Normalized eXpression (NX) levels by combining the data from the three transcriptomics datasets (HPA, GTEx and FANTOM5) using the internal normalization pipeline (*37*).

| protein | Act. | $p_{\text{tot}}$ | $t_{\text{mean}}$ | tissues of main expression |
| --- | --- | --- | --- | --- |
| PKP2 | **21** | 0.498 | 0.29 | heart muscle |
| CEP43 | 18 | 0.521 | 0.26 | testis |
| CEP135 | 14 | 0.527 | 0.30 | skeletal muscle, heart muscle, cerebral cortex, cerebellum |
| TLE5 | 13 | 0.509 | **0.24** | thymus, lymph node, testis |
| RETREG3 | 6 | 0.390 | 0.43 | prostate, thymus |
| RBM41 | 6 | 0.209 | 0.49 | pancreas, T-cells, testis, retina |
| PRIM2 | 5 | **0.536** | 0.30 | cerebellum, parathyroid gland, testis |
| MIPOL1 | 3 | 0.155 | 0.57 | pituitary gland, testis |
| REEP6 | 1 | 0.175 | 0.48 | liver, small intestine, duodenum, testis |
| HOOK1 | 1 | 0.156 | 0.46 | liver, parathyroid gland, testis, pituitary gland |
| CENPF | 1 | 0.138 | 0.46 | thymus, testis, bone marrow |
| ATP5ME | 1 | 0.096 | 0.35 | skeletal muscle, dendritic cells, heart muscle, brain |
| TRIM59 | 1 | 0.170 | 0.69 | corpus callosum, brain, T-cells, testis |
| MARK1 | 1 | N/A | >100 | epididymis, cerebral cortex, heart muscle, testis, cerebellum |
| SLC27A2 | 1 | N/A | >100 | liver, kidney |

**Tab. 2:** List of proteins targeted by SARS CoV-2 which are found in the database DisGeNet as displaying “definitive” or “strong” evidence of participating in human diseases. The Disease ID is the code of the disease in DisGeNet.

| protein | Disease ID |
| --- | --- |
| ACAD9 | C1970173 |
| AGPS | C0282529, C1838612 |
| ALG8 | C2931002, C4693472 |
| AP2M1 | C0004134, C0036572, C0557874, C0856975, C1858120, C3714756 |
| AP3B1 | C0079504, C0034069, C0024115 |
| ATP6AP1 | C4310819 |
| ATP6V1A | C4479409, C4693934 |
| BRD4 | C0018798, C0025958, C3714756, C4025871 |
| CDK5RAP2 | C1858108, C0431350 |
| CENPF | C0025958, C0152427, C0265202, C1292778 |
| CEP135 | C0431350 |
| CIT | C4310723 |
| CSDE1 | C0004352, C0557874, C3714756 |
| CYB5R3 | C0025637 |
| DNMT1 | C3807295, C1384666 |
| EXOSC3 | C1843504, C0266468 |
| GNB1 | C0557874, C0036572, C3714756, C0424605 |
| GPAA1 | C0557874 |
| IL17RA | C4310803, C3151402 |
| LARP7 | C3554439, C0013336, C0342573 |
| MOGS | C1853736 |
| MRPS2 | C4693722 |
| NDUFAF1 | C4748769, C1838979, C0751651 |
| NGLY1 | C3808991 |
| NSD2 | C0025958, C0015934, C0026827, C3714756, C0456070, C4022738 |
| PCNT | C0265202, C1292778 |
| PKP2 | C1836906, C0349788 |
| PLOD2 | C0432253, C0029434 |
| PMPCB | C4693741 |
| POLA1 | C3860213 |
| RALA | C0557874, C3714756, C0036572, C4022810 |
| RDX | C3711374, C1970239, C1384666 |
| REEP6 | C4310626, C0035334 |
| RPL36 | C1260899 |
| TAPT1 | C0029434 |
| TBK1 | C0002736, C4225325, C4693542, C0751072 |
| TCF12 | C1856266 |
| WFS1 | C0043207, C3280358, C4518338, C0011860, C1384666, C0004138 |

**Tab. 3:** Diseases identified with “definitive” or “strong” evidence in humans in the database DisGeNet for proteins targeted by SARS CoV-2. The disease ID and names are from DisGeNet.

| Disease ID | Disease/syndrome |
| --- | --- |
| C1970173 | Acyl-CoA dehydrogenase family, Member 9, Ddeficiency of |
| C0282529 | Chondrodysplasia Punctata, Rhizomelic |
| C1838612 | Rhizomelic chondrodysplasia punctata, type 3 |
| C2931002 | Congenital disorder of glycosylation type 1H |
| C4693472 | Polycystic liver disease 3 with or without kidney cysts |
| C0004134 | Ataxia |
| C0036572 | Seizures |
| C0557874 | Global developmental delay |
| C0856975 | Autistic behavior |
| C1858120 | Generalized hypotonia |
| C3714756 | Intellectual disability |
| C0079504 | Hermanski-Pudlak syndrome |
| C0034069 | Pulmonary fibrosis |
| C0024115 | Lung diseases |
| C4310819 | Immunodeficiency 47 |
| C4479409 | Cutis laxa, autosomal recessive, type IID |
| C4693934 | Epileptic encephalopathy, infantile or early childhood, 3 |
| C0018798 | Congenital heart defects |
| C0025958 | Microcephaly |
| C4025871 | Abnormality of the face |
| C1858108 | Microcephaly, primary autosomal recessive, 3 |
| C0431350 | Primary microcephaly |
| C0152427 | Polydactyly |
| C0265202 | Seckel syndrome |
| C1292778 | Chronic myeloproliferative disorder |
| C4310723 | Microcephaly 17, primary, autosomal recessive |
| C0004352 | Autistic disorder |
| C0025637 | Methemoglobinemia |
| C3807295 | Cerebellar ataxia, deafness, and narcolepsy, autoomal dominant |
| C1384666 | Hearing impairment |
| C1843504 | Pontocerebellar Hhypoplasia Ttype 1 |
| C0266468 | Congenital pontocerebellar hypoplasia |
| C0424605 | Developmental delay (disorder) |
| C4310803 | Immunodeficiency 51 |
| C3151402 | Candidiasis, familial, 5 |
| C3554439 | Microcephalic primordial dwarfism Alazami type |
| C0013336 | Dwarfism |
| C0342573 | Pituitary dwarfism I |
| C1853736 | Congenital disorder of glycosylation, Type IIB |
| C4693722 | Combined oxydative phosphorylation deficiency 36 |
| C4748769 | Mitochondrial complex I deficiency, nuclear type 11 |
| C1838979 | Mitochondrial complex I deficiency |
| C0751651 | Mitochondrial diseases |
| C3808991 | NGLY1 deficiency |
| C0015934 | Fetal Ggrowth retardation |
| C0026827 | Muscle hypotonia |
| C0456070 | Growth delay |
| C4022738 | Neurodevelopmental delay |
| C1836906 | Arrhythmogenic right ventricular dysplasia, familial, 9 |
| C0349788 | Arrhythmogenic right Vventricular dysplasia |
| C0432253 | Bruck syndrome |
| C0029434 | Osteogenesis imperfecta |
| C4693741 | Multiple mitochondrial dysfunctions syndrome 6 |
| C3860213 | Autoinflammatory disorder |
| C4022810 | Abnormality of nervous system morphology |
| C3711374 | Nonsyndromic deafness |
| C1970239 | Deafness, autosomal recessive, 24 |
| C4310626 | Retinitis pigmentosa 77 |
| C0035334 | Retinitis Pigmentosa |
| C1260899 | Anemia, Diamond-Blackfan |
| C0002736 | Amyotrophic Lateral Sclerosis |
| C4225325 | Frontotemporal dementia and/or amyotrophic lateral sclerosis 4 |
| C4693542 | Encephalopathy, acute, infection-induced (Herpes-specific), Susceptibility to, 8 |
| C0751072 | Frontotemporal lobar degeneration |
| C1856266 | Coronal craniosynostosis |
| C0043207 | Wolfram syndrome |
| C3280358 | Wolfram-like syndrome, autosomal dominant |
| C4518338 | Wolfram-like syndrome |
| C0011860 | Diabetes Mmellitus, non-Insulin-dependent |
| C0004138 | Ataxias, hereditary |

**Tab. 4:**
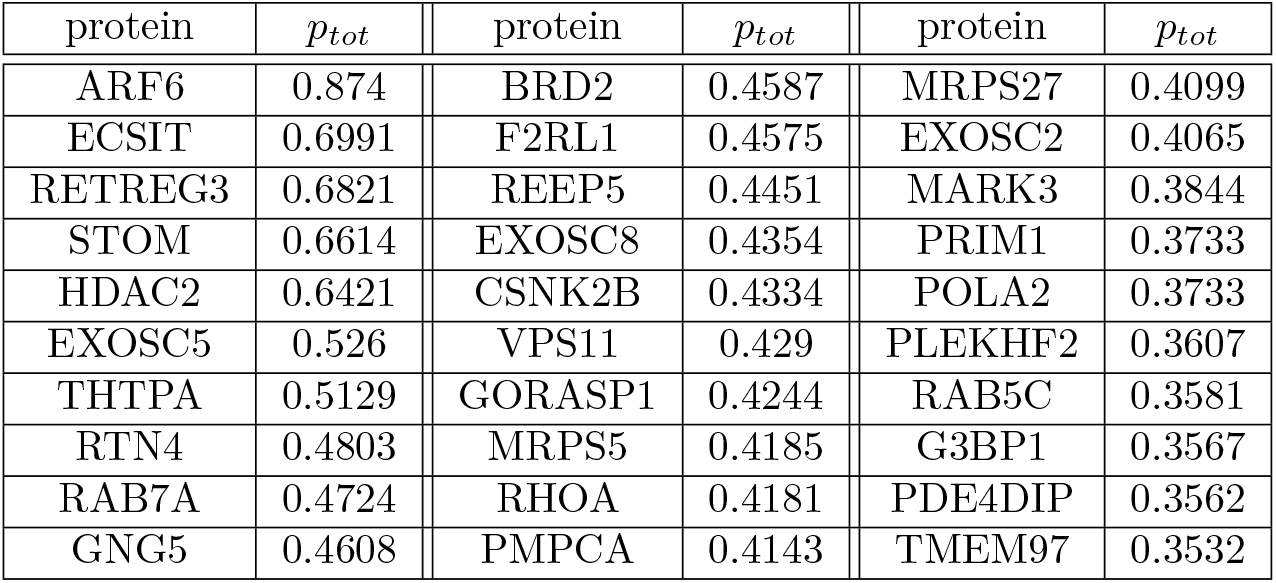
Proteins with the largest effect on perturbing those 38 proteins identified in human diseases. *p*_tot_ is the sum of the probabilities that the given protein activates those identified as having “definitive” or “strong” evidence of being involved in a human disease.

**Tab. 5:** RNA expression overview for proteins targeted by SARS CoV-2 and mainly expressed outside the lungs. We select the top RNA expressions in the four databases reported in The Human Protein Atlas.

| protein | consensus | HPA | GTEx | FANTOM5 |
| --- | --- | --- | --- | --- |
| PKP2 | heart muscle | heart muscle<br>fallopian tube | heart muscle | heart muscle |
| CEP43 | testis | testis | testis | testis |
| CEP135 | thymus | lymph node<br>endometrium<br>testis | testis | thymus |
| TLE5 | skeletal muscle<br>heart muscle | cerebral cortex | cerebellum | cerebellum<br>heart muscle |
| RETREG3 | pancreas | T-cells<br>testis | testis | retina<br>testis<br>endometrium |
| RBM41 | cerebellum<br>parathyroid<br>gland | parathyroid<br>gland | testis<br>thyroid gland | cerebellum |
| PRIM2 | prostate<br>thymus | prostate | N/A | thymus |
| MIPOL1 | pituitary gland | testis | testis | pituitary gland |
| REEP6 | liver<br>small intestine | duodenum<br>small intestine<br>testis | testis | testis<br>small intestine |
| HOOK1 | liver | parathyroid<br>gland | testis | pituitary gland<br>kidney |
| CENPF | thymus | testis<br>bone marrow | testis | thymus |
| ATP5ME | skeletal muscle | dendritic cells | heart muscle<br>brain | heart muscle<br>spinal cord<br>mid brain |
| TRIM59 | corpus callosum<br>mid brain<br>thalamus<br>thymus | granulocytes<br>T-cells | testis<br>spinal cord | thalamus<br>mid brain<br>corpus callosum |
| MARK1 | epididymis<br>cerebral cortex<br>heart muscle | cerebral cortex<br>testis<br>epididymis | cerebellum | cerebellum |
| SLC27A2 | liver | kidney<br>liver | liver | liver |

**Tab. 6:** List of proteins mainly expressed outside the lungs and their major activators, which are proteins mainly expressed in the lungs.

| protein | main perturbors from the lungs |
| --- | --- |
| PKP2 | ARF6, RHOA, CSNK2A2, CSNK2B, GOLGA2, PRKACA, PRKAR2B, RAB1A, RAP1GDS1, BRD2, TLE1, NUP214, GPAA1, PMPCB, PDE4DIP, AKAP9, SUN2, NGDN, GRIPAP1, THTPA, ALG8 |
| CEP43 | PCNT, PRKAR2A, PRKAR2B, RAB7A, PDE4DIP, CEP350, AKAP9, TUBGCP3, TUBGCP2, CNTRL, CEP250, NINL, CEP68, PLEKHA5, CDK5RAP2, MIB1, GCC1, TIMM29 |
| CEP135 | GOLGB1, HDAC2, RAB8A, PRKAR2A, PRKAR2B, RAB1A, PDE4DIP, CEP350, ERC1, IL17RA, ZNF318, TBK1, PLEKHA5, CDK5RAP2, MIB1 |
| TLE5 | GOLGA2, TLE1, TLE3, EIF4E2, NUTF2, EXOSC8, IL17RA, EXOSC3, UBAP2, EXOSC5, GCC1, ZNF503, USP54, EXOSC2 |
| RETREG3 | STOM, REEP5, NDUFAF1, RTN4, PLEKHF2, CYB5B |
| RBM41 | GOLGA2, RAB2A, TUBGCP2, NINL, GORASP1, GCC1 |
| PRIM2 | POLA1, PRIM1, RAE1, ERP44, POLA2 |
| MIPOL1 | EIF4E2, IL17RA, USP54 |
| REEP6 | REEP5 |
| HOOK1 | VPS39 |
| CENPF | G3BP1 |
| ATP5ME | SRP72 |
| TRIM59 | ECSIT |
| MARK1 | MARK2 |
| SLC27A2 | USP13 |

### 3.4 Subdiffusion paths

Finally we study here how the diffusive process determines the paths that the perturbation follows when diffusing from a protein to another not directly connected to it. The most efficient way of propagating a perturbation between the nodes of a network is through the shortest (topological) paths that connect them. The problem for a (sub)diffusive perturbation propagating between the nodes of a network is that it does not have complete information about the topology of the network as to know its shortest (topological) paths. The network formed by the proteins targeted by SARS CoV-2 is very sparse and this indeed facilitates that the perturbations occurs by following the shortest (topological) paths most of the time. Think for instance in a tree, which has the lowest possible edge density among all connected networks. In this case, the perturbation will always use the shortest (topological) paths connecting pairs of nodes. However, in the case of the PPI network studied here a normal diffusive process, i.e., *α* =1, not always uses the shortest (topological) paths. In this case, there are 1294 pairs of proteins for which the diffusive particle uses a shortest diffusive path which is one edge longer than the corresponding shortest (topological) path. This represents 6.11% of all total pairs of proteins that are interconnected by a path in the PPI network of proteins targeted by SARS CoV-2. However, when we have a subdiffusive process, i.e., *α* = 0.75, this number is reduced to 437, which represents only 2.06% of all pairs of proteins. Therefore, the subdiffusion process studied here through the PPI network of proteins targeted by SARS CoV-2 has an efficiency of 97.9% relative to a process that always uses the shortest (topological) paths in hopping between proteins. In Fig. 5 we illustrate the frequency with which proteins not in the shortest (topological) paths are perturbed as a consequence that they are in the shortest subdiffusive paths between other proteins. For instance, the following is a shortest diffusive path between the two endpoints: RHOA-PRKACA-PRKAR2A-CEP43-RAB7A-ATP6AP1. The corresponding shortest (topological) path is: RHOA-MARK2-AP2M1-RAB7A-ATP6AP1, which is one edge smaller. The proteins PRKACA, PRKAR2A and CEP43 are those in the diffusive path which are not in the topological one. Repeating this selection process for all the diffusive paths that differs from the topological ones we obtained the results illustrated in Fig. 5. As can be seen there are 36 proteins visited by the shortest diffusive paths which are not visited by the corresponding topological ones. The average degree of these proteins is 7.28 and there is only a small positive trend between the degree of the proteins and the frequency with which they appear in these paths, e.g., the Pearson correlation coefficient is 0.46.

**Fig. 5:**
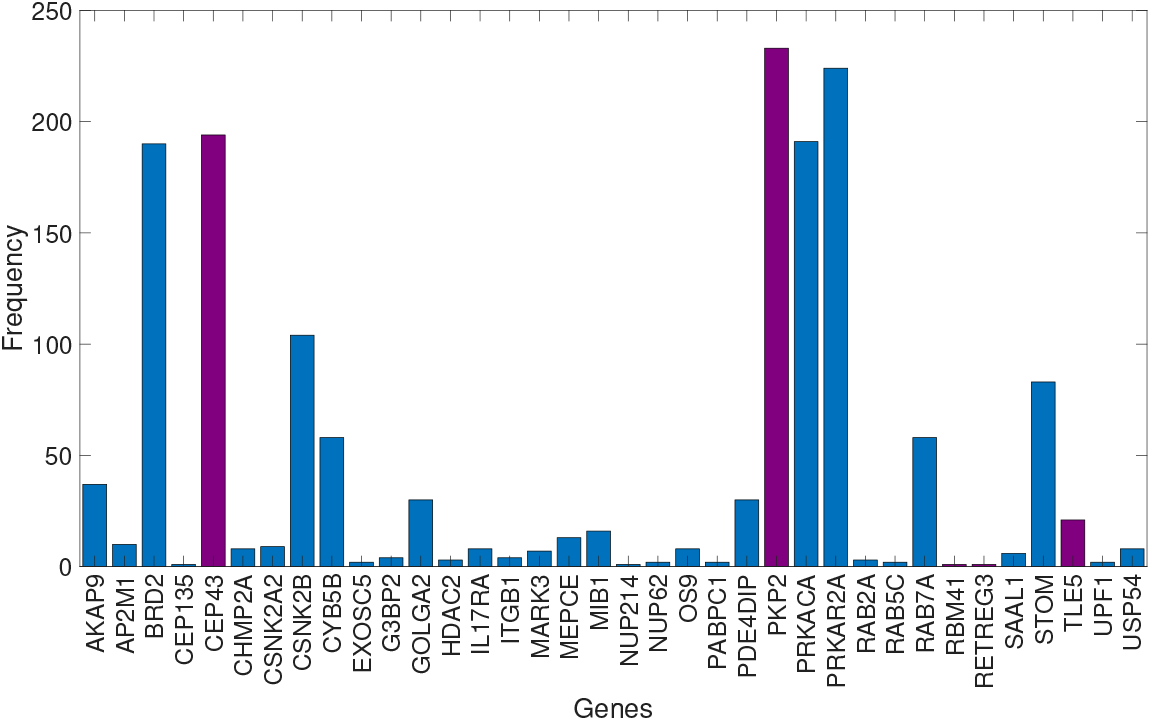
Proteins in shortest subdiffusive paths. Frequency of proteins that appear in the shortest subdiffusion paths but not in the shortest (topological) paths connecting any pair of nodes in the PPI network. Bars marked in wine color are for those proteins expressed mainly outside the lungs.

## 4 Discussion

We have presented a methodology that allow the study of diffusive processes in (PPI) networks varying from normal to subdiffusive regimes. Here we have studied the particular case in which the time-fractional diffusion equation produces a subdiffusive regime, with the use of *α* = 3/4 in the network of human proteins targeted by SARS CoV-2. A characteristic feature of this PPI network is that the second smallest eigenvalue is very small, i.e., *μ*_2_ = 0.0647. As this eigenvalue determines the rate of convergence to the steady state, the subdiffusive process converges very slowly to that state. What it has been surprising is that even in these conditions of very small convergence to the steady state, there is a very early increase of the probability in those proteins closely connected to the initiator of the diffusive process. That is, in a subdiffusive process on a network, the time at which a perturbation is transmitted from the initiator to any of its nearest neighbors occurs at an earlier time than for the normal diffusion. This is a consequence of the fact that *E*_*α*,1_ (− *γt^α^*) decreases very fast at small values of *t^α^*, which implies that the perturbation occurring at a protein *i* at *t* = 0 is transmitted almost instantaneously to the proteins closely connected to *i*. This effect may be responsible for the explanation about why subdiffusive processes, which are so globally slow, can carry out cellular processes at a significant rate in cells. We have considered here a mechanism consisting in switching and restarting several times during the global cellular process. For instance, a subdiffusive process starting at the protein *i* perturbs its nearest neighbors at very early times, among which we can find the protein *j*. Then, a new subdiffusive process can be restarted again at the node *j*, and so on.

One of the important findings of using the current model for the study of the PIN of proteins affected by SARS CoV-2 is the identification of those proteins which are expressed outside the lungs that can be more efficiently perturbed by those expressed in the lungs (see 1). For instance, the protein with the largest number of activators, PKP2, appears mainly in the heart muscle. It has been observed that the elevation of cardiac biomarkers is a prominent feature of COVID-19, which in general is associated with a worse prognosis (*41*). Myocardial damage and heart failure are responsible for 40% of death in the Wuhan cohort (see references in (*41*)). Although the exact mechanism involving the heart injury is not known the hypothesis of direct myocardial infection by SARS CoV-2 is a possibility, which act along or in combination with the increased cardiac stress due to respiratory failure and hypoxemia, and or with indirect injury from the systemic inflammatory response (*41, 42, 43, 44*). As can be seen in n Table 1 testis is the tissue where several of the proteins targeted by SARS CoV-2 are mainly expressed, e.g., CEP43, TLE5, PRIM2, MIPOL1, REEP6, HOOK1, CENPF, TRIM59, and MARK1. Currently there is not conclusive evidence about testis damage by SARS CoV-2 (*45, 46, 47, 48*). However, the previous SARS CoV that appeared in 2003 and which shares 82% of proteins with the current one, produced testis damage and spermatogenesis, and it was concluded that orchitis was a complication of that previous SARS disease (*45*). We also detect a few proteins mainly expressed in different brain tissues, such as CEP135, PRIM2, TRIM59, and MARK1. The implication of SARS CoV-2 and cerebrovascular diseases has been reported, including neurological manifestations as well as cerebrovascular disease, such as ischemic stroke, cerebral venous thrombosis, and cerebral hemorrhage (*49, 50, 51*).

Kidney damage in SARS CoV-2 patients has been reported (*52, 53, 54*), which includes signs of kidney dysfunctions, proteinuria, hematuria, increased levels of blood urea nitrogen, and increased levels of serum creatinine. As much as 25% of acute kidney injury has been reported in clinical setting of SARS CoV-2 patients. One of the potential mechanisms for kidney damage is the organ crosstalk (*52*), as can be the mechanism of diffusion from proteins in the lungs to proteins in the urinary tract and kidney proposed here. A very interesting observation from Table 1 is the existence of several proteins expressed mainly in the thymus and T-cells, such as: TLE5, RETREG3, RBM41, CENPF, and TRIM59. It has been reported that many of the patients affected by SARS CoV-2 in Wuhan displayed significant decrease of T-cells (*55*). Thymus is an organ which displays a progressive decline with age with reduction of the order of 3-5% a year until approximately 30-40 years of age and of about 1% per year after that age. Consequently, it was proposed that the role of thymus should be taken into account in order to explain why COVID-19 appears to be so mild in children (*55*). The protein TLE5 is also expressed significantly in the lymph nodes. It was found by Feng et al. (*56*) that SARS CoV-2 induces lymph follicle depletion, splenic nodule atrophy, histiocyte hyperplasia and lymphocyte reductions. The proteins HOOK1 and MIPOL1 are significantly expressed in the pituitary gland. There has been some evidence and concerns that COVID-19 may also damage the hypothalamo-pituitary-adrenal axis has been expressed by Pal (*57*), which may be connected with the participation of the before mentioned proteins.

Another surprising finding of the current work is the elevated number of subdiffusive shortest paths that coincide with the shortest (topological) paths connecting pairs of proteins in the PPI of human proteins targeted by SARS CoV-2. This means that the efficiency of the diffusive paths connecting pairs of nodes in this PPI is almost 98% in relation to a hypothetical process which uses the shortest (topological) paths in propagating perturbations between pairs of proteins. The 437 shortest diffusive paths reported here contain one more edge than the corresponding shortest (topological) paths. The proteins appearing in these paths would never be visited in the paths connecting two other proteins if only the shortest (topological) paths were used. What it is interesting is to note that 6 out of the 15 proteins which are mainly expressed outside the lungs are among the ones “crossed” by these paths. They are: TLE5 (thymus, lymph node, testis), PKP2 (heart muscle), CEP135 (skeletal muscle, heart muscle, cerebral cortex, cerebellum), CEP43 (testis), RBM41 (pancreas, T-cells, testis, retina) and RETREG3 (prostate, thymus). This means that the perturbation of these proteins occurs not only through the diffusion from other proteins in the lungs directly to them, but also through some “accidental” diffusive paths between pairs of proteins which are both located in the lungs.

All in all, the use of time-fractional diffusive models to study the propagation of perturbations on PPI networks seems a very promising approach. The model is not only biologically sounded but it also allows us to discover interesting hidden patterns of the interactions between proteins and the propagation of perturbations among them. In the case of the PIN of human proteins targeted by SARS CoV-2 our current finding may help to understand potential molecular mechanisms for the multi-organs and systemic failures occurring in many patients.

## Acknowledgements

The author thanks Dr. Deisy Morselli Gysi for sharing data and information.

## Competing interests

The author declares that he has no competing interests.

## 5 Supplementary Information

### Supplementary Note 1

We first prove that the solution of the FTD equation is given by 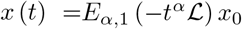, where *E*_*α*,1_ (*M*) is the Mittag-Leffler matrix function of *M*. We use the spectral decomposition of the network Laplacian 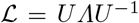, where 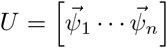 and *Λ* = *diag* (*μ_r_*). Then we can write

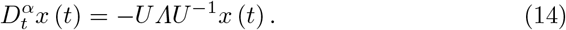

Let us define *y* (*t*) = *U*^−1^*x* (*t*), such that 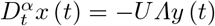 and we have

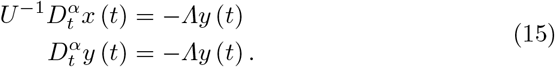

As Λ is a diagonal matrix we can write

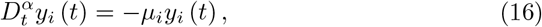

which has the solution

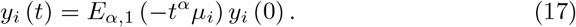

We can replace *y_i_* (*t*) = *U*^−1^*x_i_* (*t*) to have

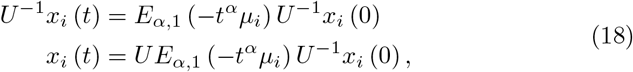

which finally gives the result in matrix-vector when written for all the nodes,

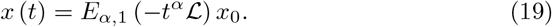

### Supplementary Note 2

In this Note we prove that the function 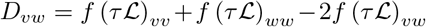, where 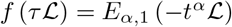 is a Euclidean distance between the corresponding pair of nodes in the network. The matrix function 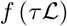 can be written as 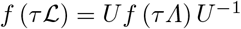. Let 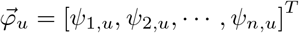. Then,

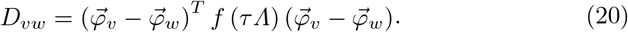

Therefore, because 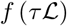 is positive defined we can write

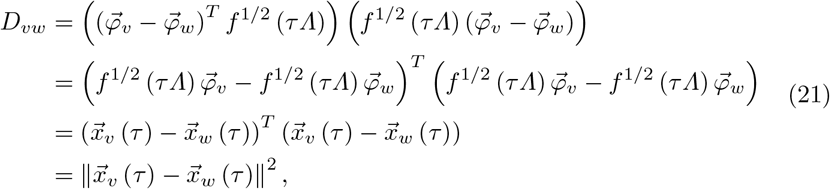

where 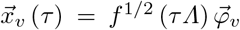. Consequently, *D_vw_* is a square Euclidean distance between *v* and *w*. In this sense the vector 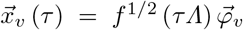 is the position vector of the node *v* in the diffusion Euclidean space. Because 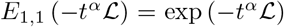 we have that *D_vw_* generalizes the diffusion distance studied by Coifman and Lafon, which is the particular case when *α* =1.

## Notes

### Competing Interest Statement

The authors have declared no competing interest.

